# An insect’s view of a numerosity illusion: a simple strategy may explain complex numerical performance in bumblebees

**DOI:** 10.1101/2023.08.22.554303

**Authors:** Elia Gatto, Cui Guan, Agrillo Christian, Simone Cutini, Lars Chittka, Maria Elena Miletto Petrazzini

## Abstract

Despite some similarities in numerical abilities among vertebrates, animals seem to differ with respect to the perceptual factors underlying numerical estimation. This is particularly evident in the case of numerosity illusions, a type of illusory phenomena elicited by the spatial arrangement of the elements that compose it. We investigated whether an insect experiences the most famous numerosity illusion, the Solitaire Illusion. Bumblebees (*Bombus terrestris audax*) presented with two mixed arrays containing magenta and yellow dots were trained to select the array containing the larger number of a target colour (either magenta or yellow). In the test phases, bumblebees were presented with sets of stimuli differing in the numerical ratio, including the illusory arrays: in one array, magenta dots were centrally located (yellow dots in the perimeter), whereas in the other one, magenta dots were located in the perimeter (yellow dots in the centre). Bees discriminated up to a ratio of 0.78 (14 vs. 18). Video analyses showed that bumblebees sequentially scanned the stimuli before making a choice. Bumblebees required longer scanning time for patterns with smaller numbers of reinforced colour dots, suggesting that bumblebees spent longer time on searching for more information or finding a reinforced arrangement of spatial dots before making an accurate decision. In the presence of the “illusion arrangement”, bees behaved in a manner consistent with misperception of numerosity, overestimating the number of centrally located dots. The spatial configuration of the stimuli influenced how the patterns were assessed: bees examined the clustered reinforced colour dots longer. Flying paths analysis suggested that bumblebees followed the paths formed by similar colour dots rather than counting each dot to make a numerical discrimination. These results challenge the notion that visual illusions can be inferred simply from evaluating choices of different visual patterns and reinforce the idea that multiple strategies can potentially be used to achieve seemingly similar cognitive outcomes. Detailed analyses of decision-making processes are essential to deduce the cognitive strategies underpinning discrimination tasks, whether they are numerical or other visual tasks.

## Introduction

While mathematical abilities have been historically regarded as being a uniquely human trait, it is now clear that many animal species possess a predisposition to use quantity information without symbolic representation [1–4]. This capacity appears to play a significant role in several ecological contexts, such as foraging, mate choice, and social interaction [5–8]. Recent studies, however, have questioned the existence of identical perceptual/cognitive mechanisms between humans and their close relatives. These studies investigated the role of item clustering in numerical estimation in humans, apes, and monkeys by using a well-known numerosity illusion as a research tool, the Solitaire Illusion [9–10]. This illusion occurs when we misperceive the relative number of two different colours of otherwise identical objects, depending on their spatial arrangements. The illusion is robust in humans and can be consistently observed in different experimental contexts [9, 11–13]. For understanding the susceptibility to this illusion, it is necessary to delve into the mechanism underlying its perceptual mechanism. The misperception of numerosity estimation in this illusion is triggered by the spatial configuration of elements [9]. According to the Gestalt principles [14], a set of same-size items lead to be visually grouped in a better figure than a set of varied-size items (similarity principles). In the same way, closed-spaced items form a better figure than spared items (proximity principles). Taken together, sets of elements which form a good Gestalt, i.e., set of similar and close-related items, have been proposed to appear more numerous [9, 15] thus the spatial configuration of coloured dots rather than the pure numerical information is used to make numerical judgement.

In recent years, research on the cognitive system of invertebrate organisms (such as bees), has raised growing interest among comparative psychologists. Bees are characterized by impressive cognitive capacities, including the formation of concepts such as sameness and difference [16, 17], object manipulation skills learned by observing conspecifics [18, 19], and even numerical abilities [20–22]. However, there might be profound differences in the visual-behavioural strategies by which insects and vertebrates solve numerosity tasks [22], and therefore an examination of the Solitaire Illusion in these animals is of interest, as are the choices processes leading to decisions in such a task. In visual processing, primates, other mammals, and birds can solve visual discrimination at glance, as numerical estimation and visual categorization [23]. Large brain size improves the parallel integration of different sensory information and an increases representational capacity of the objects, rather than necessarily affecting cognitive capacity [24]. The nature of parallel visual processing in vertebrates may exacerbate the misperception of the illusion. Another mechanism observed in several species rely more predominantly on acquiring visual information by actively sampling their environment, a mechanism defined as “active vision” [25, 26]. Such mechanisms might be evolved to compensate a less capacity of processing information from several visual sources at the same time, and consequently, with a continuous dependence on systematically scanning the scene [24, 27]. Indeed, individuals typically require scanning stimuli from close by [28, 29], inspecting the elements that compose the scene one by one [8, 22] and can not discriminate between complex stimuli when presented for less than 100ms [28]. Such visual behaviours compensate for the lack of parallel processing a whole image at a glance [24].

Here we addressed how an invertebrate species, *Bombus terrestris audax* responds to patterns that induce a numerosity illusion in humans, the Solitaire illusion. We also explored the tactics of pattern inspection applied by bees solving the numerical discrimination.

## Results

We individually trained naïve bumblebees (*Bombus terrestris audax*, n = 20; 3 colonies) to discriminate between pairs of mixed arrays of magenta and yellow dots arranged in a cross-shape pattern on a white background vertically presented in a flight arena. Each array consisted of 32 dots but the ratio between the number of magenta and yellow dots in the two arrays varied across the experimental phases. During the training phase, we used two sets of stimuli: one with 11 vs. 21 dots (ratio 0.52) and one with 13 vs. 19 dots (ratio 0.68). We trained one group of bumblebees (*n* = 10) to select the array with the larger number of magenta dots, and a second group (*n* = 10) was trained to select the array with the larger number of yellow dots. We positively reinforced correct choices with 10 μL 50% sucrose solution and 10 μL saturated quinine hemisulfate solution was used as punishment for incorrect choices.

During the training phase, both groups of bees learned the task and significantly chose the patterns with the larger numbers of dots (overall performance 73% ± 10%, *t*_19_ = 10.01, *P* < 0.01, *Cohen’s d* = 2.24) across the training sessions (ß = 4.22, C.I. [2.43, 6.02], F_1,76.34_ = 28.53, *P* < 0.01, ηp^2^ = 0.27). No difference was found between the two groups trained to yellow and magenta dots (ß = 9.13, C.I. [−2.41, 20.67], F_1,49.30_ = 2.32, *P* = 0.13, ηp^2^ = 0.05), nor was the interaction between the reinforced colour and training significant (ß = −1.52, C.I. [−4.06, 1.01], F_1,76.34_ = 1.38, *P* = 0.24, ηp^2^ = 0.02; Fig 2A).

**Figure 1.**
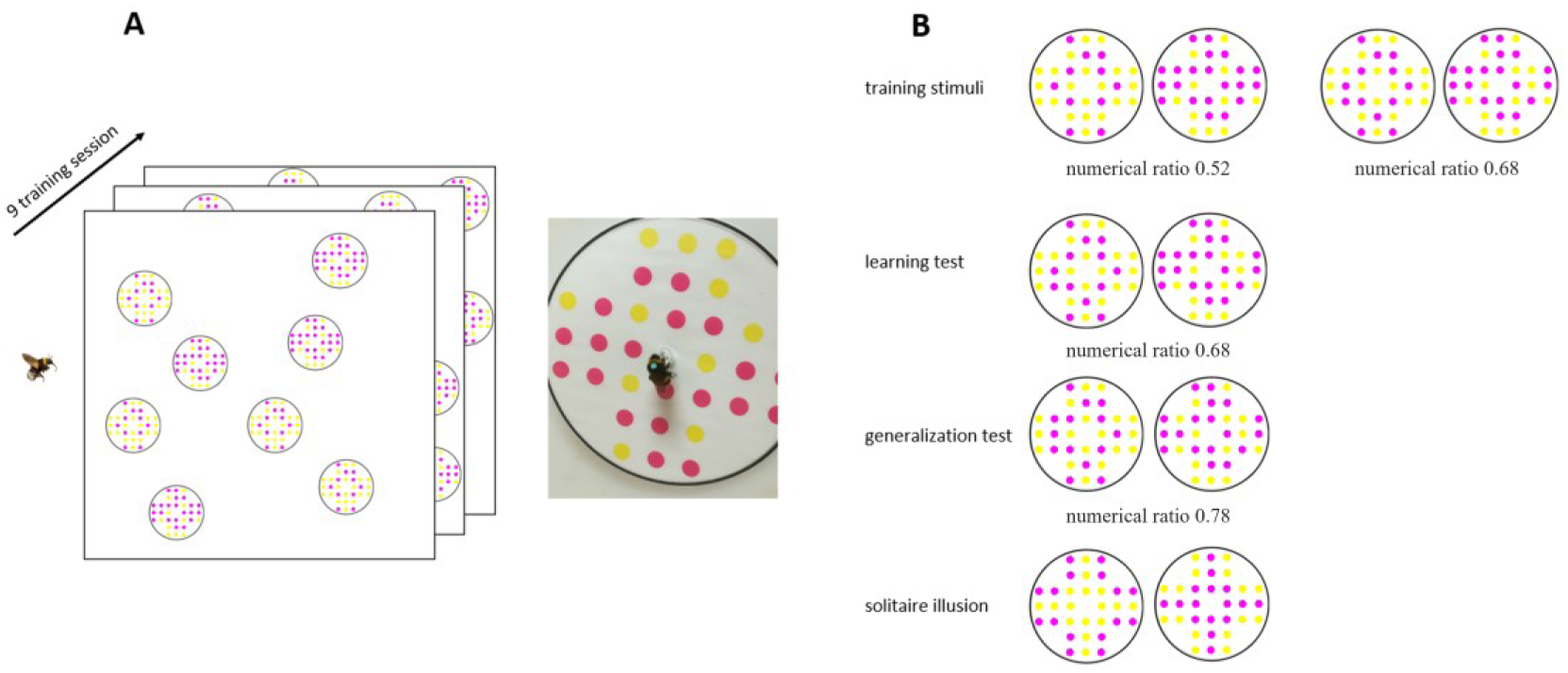
Training protocol and stimuli. (**A**) Each subject was trained in 9 consecutive trials, in which four pairs of the same numerical comparison (**B**) were presented. Training and test patterns were 12 cm diameter disks with 32 dots differing in colour (yellow and magenta) and numerosity (0.50 and 0.68 ratio in training phase; 0.68, 0.78, and 1 in test phase). Each pattern was attached via its centre to the vertical wall of the flight arena by a plastic tip (5 mm diameter). The distribution of patterns was randomized during the experiment. In each trial, subjects were free to forage and return to the hive. Following the 9 training trials, bees were subjected to three non-reinforced test trials (sterile distilled water was provided). Responses were analysed from video recording of the first 120 s in the flight arena.

**Figure 2.**
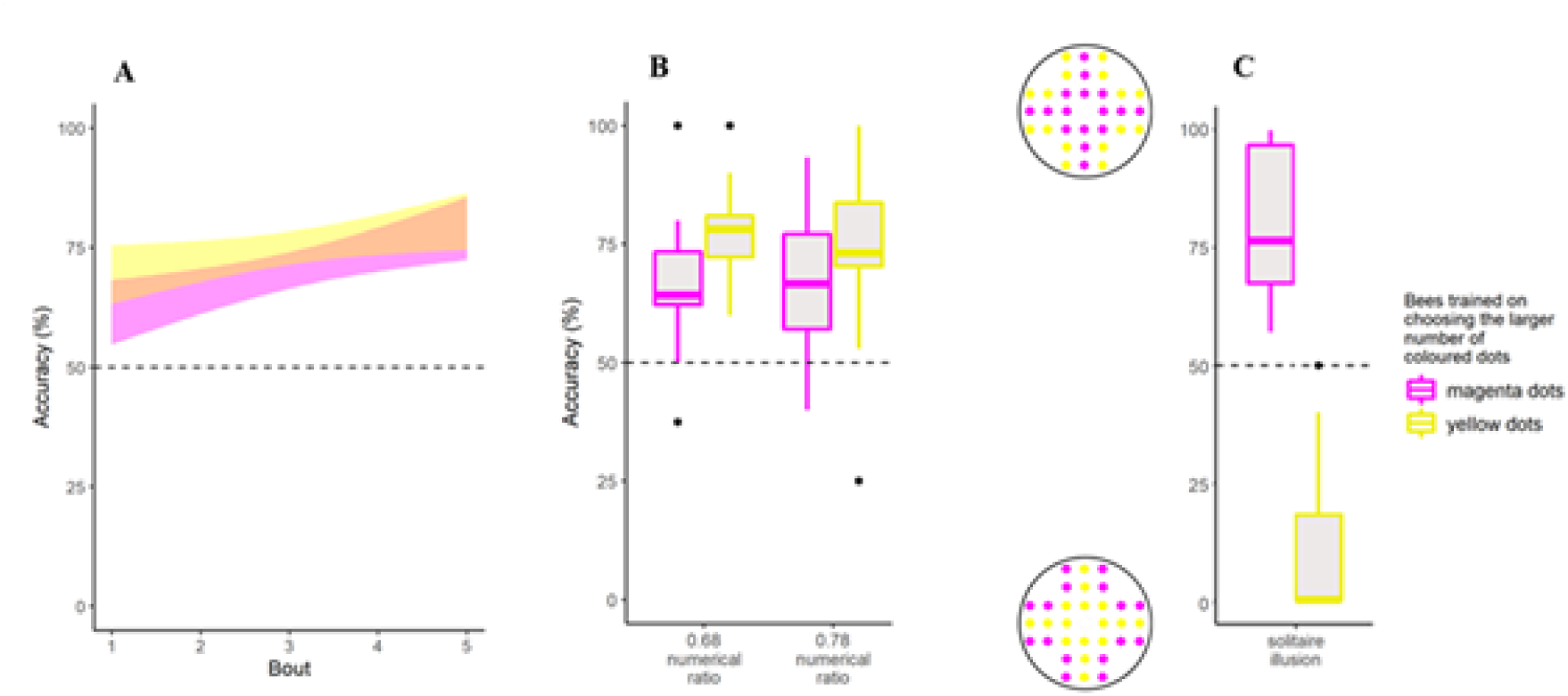
Learning curves during the training phase and performance during the test phases. (**A**) A generalised linear mixed-effects model (see methods) shows a significant increase of performance during the last 50 choices (5 blocks of 10 choices each) of the training phase (*P* < 0.001), irrespective of the positively reinforced pattern (*P* > 0.279). (**B**) The median percentage (±SD) of correct choices is plotted as a function of tests (learning test and transfer test) and the reinforced patterns. The black dashed line indicates the chance level performance (50%). Overall, bees chose accurately the correct stimulus in the learning (*P* < 0.01) and in the generalization test (*P* < 0.01). (**C**) Percentage of individual choices in the presence of the “solitaire illusion”. The Y-axis refers to the percentage of choices towards the clustered or spared configuration of reinforced colour dots, while the dotted line represented the equally number of responses to the two patterns. Overall, bees significantly choose the stimuli in which the reinforced colour is spatially clustered (*P* < 0.01). There is no difference as a function of the reinforced colour.

In the learning test, bumblebees chose the array associated with the larger number of reinforced colour dots (overall performance 72% ± 15%, *Estimate* = 0.93, *SD* = 0.15, *z* = 6.34, *P* < 0.01; effect of positively reinforced pattern: ß = 0.49, C.I. [−0.10, 1.10], χ^2^_1_ = 2.67, *P* = 0.10, *delta* - *R*^2^ = 0.01; Figure 2B). The same pattern was observed in the transfer test (overall performance 70% ± 19%, *Estimate* = 0.94, *SD* = 0.22, *z* = 4.29, *P* < 0.01; effect of positively reinforced pattern: ß = 0.39, C.I. [−0.47, 1.29], χ^2^_1_ = 0.87, *P* = 0.35, *delta* - *R*^2^ = 0.07; Figure 2B). There was no difference between the two numerical ratios (*t*_19_ = 0.46, *P* = 0.65, *Cohen’s D* = 0.10).

The video-analysis of scanning behaviour in the learning test suggested that bumblebees’ decision speed is influenced by the nature of information. Indeed, bumblebees on average scanned the stimulus with smaller number of reinforced colour dots for longer periods (*F*_1,437.16_ = 4.19, *P* = 0.04, ηp^2^ = 0.01), while no significant difference was found between the positively reinforced colour dots (*F*_1, 17.97_ = 0.01, *P* = 0.94, ηp^2^ = 0.00), nor the interaction between the factors (*F*_1,437.16_ = 0.61, *P* = 0.44, ηp^2^ = 0.00), indicating that in this particular arrangement of visual stimuli, bumblebees required longer sampling time to obtain information for making an accurate decision especially when the information is scarce (i.e., stimulus with smaller number of reinforced colour dots). Bumblebees scanned the bottom and middle areas of the pattern more than the top area (bottom | middle: *Estimate* = 0.01, *SD* = 0.12, *z* = 0.05, 95% CI [−0.22, 0.24]; middle|top: *Estimate* = 2.14, *SD* = 0.13, *z* = 16.16, 95% CI [1.88, 2.40]; Figure 3). No significant difference was found between the types of patterns (ß = 0.17, C.I. [0.00, 0.62], χ^2^_1_ = 1.53, *P* = 0.22), or between the positively rewarded colour dots (ß = 0.31, C.I. [−0.10, 0.44], χ^2^_1_ = 0.11, *P* = 0.74), but the interaction between these two factors was significant (ß = −0.52, C.I. [−0.89, - 0.15], χ^2^_1_ = 7.70, *P* < 0.01; *McFadden’s - R*^2^ < 0.01).

**Figure 3.**
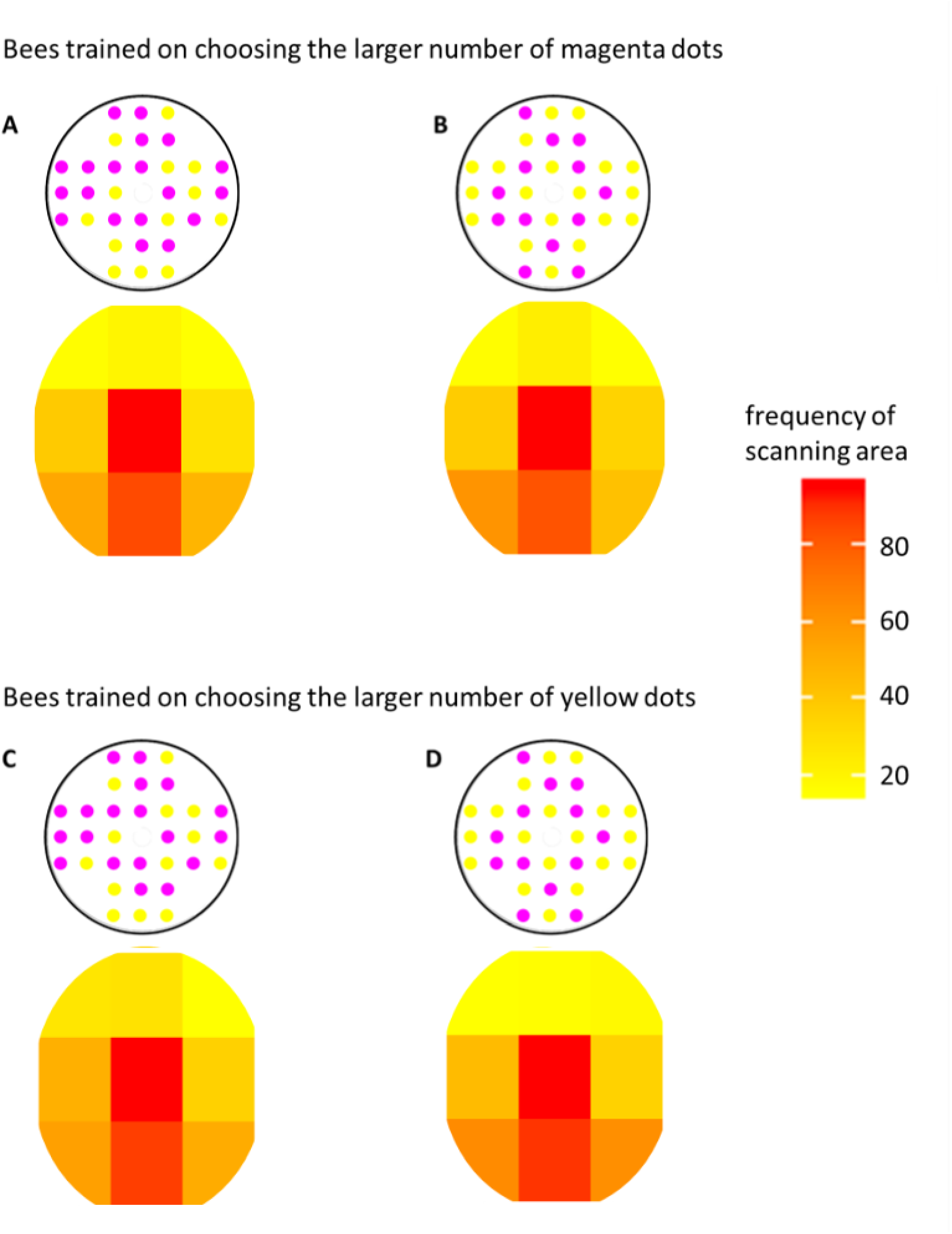
Scanning behaviour of bees in the presence of the learning test. Frequency maps of bee location in front of a pattern during solitaire illusion tests. Bees spent significantly more time on scanning the lower and central areas of each pattern (bottom|middle: 95% CI [−0.22, −0.24]; middle|top: 95% CI [1.88, 2.40]).

A similar result was found when analysing the scanning behaviour of bees during the transfer test. Bumblebees scanned the stimulus with smaller number of reinforced colour dots longer (*F*_1,449.42_ = 4.25, *P* = 0.04, ηp^2^ = 0.01). No other factor was significant (Ps > 0.21), suggesting that, in a challenging discrimination, bumblebees increase sampling time to gather high-quality information and make properly decision. Again, bumblebees scanned the bottom and middle area of the pattern more than the top area (bottom|middle: *Estimate* = - 0.26, *SD* = 0.13, *z* = - 2.10, 95% CI [−0.51, −0.02]; middle|top: *Estimate* = 1.79, *SD* = 0.13, *z* = 13.33, 95% CI [1.53, 2.06]; Figure 4). A significant difference was found between the types of stimuli (ß = −0.17, C.I. [−0.43, 0.08], χ^2^ = 6.06, *P* = 0.01). The analysis did not reveal a significant effect of rewarded colour dots (ß = −0.04, C.I. [−0.38, 0.30], χ^2^ = 0.32, *P* = 0.57), nor the interaction stimulus × reinforced colour dots (ß = −0.09, C.I. [−0.45, 0.26], χ^2^ = 0.26, *P* = 0.61, *McFadden’s - R*^2^ < 0.01).

**Figure 4.**
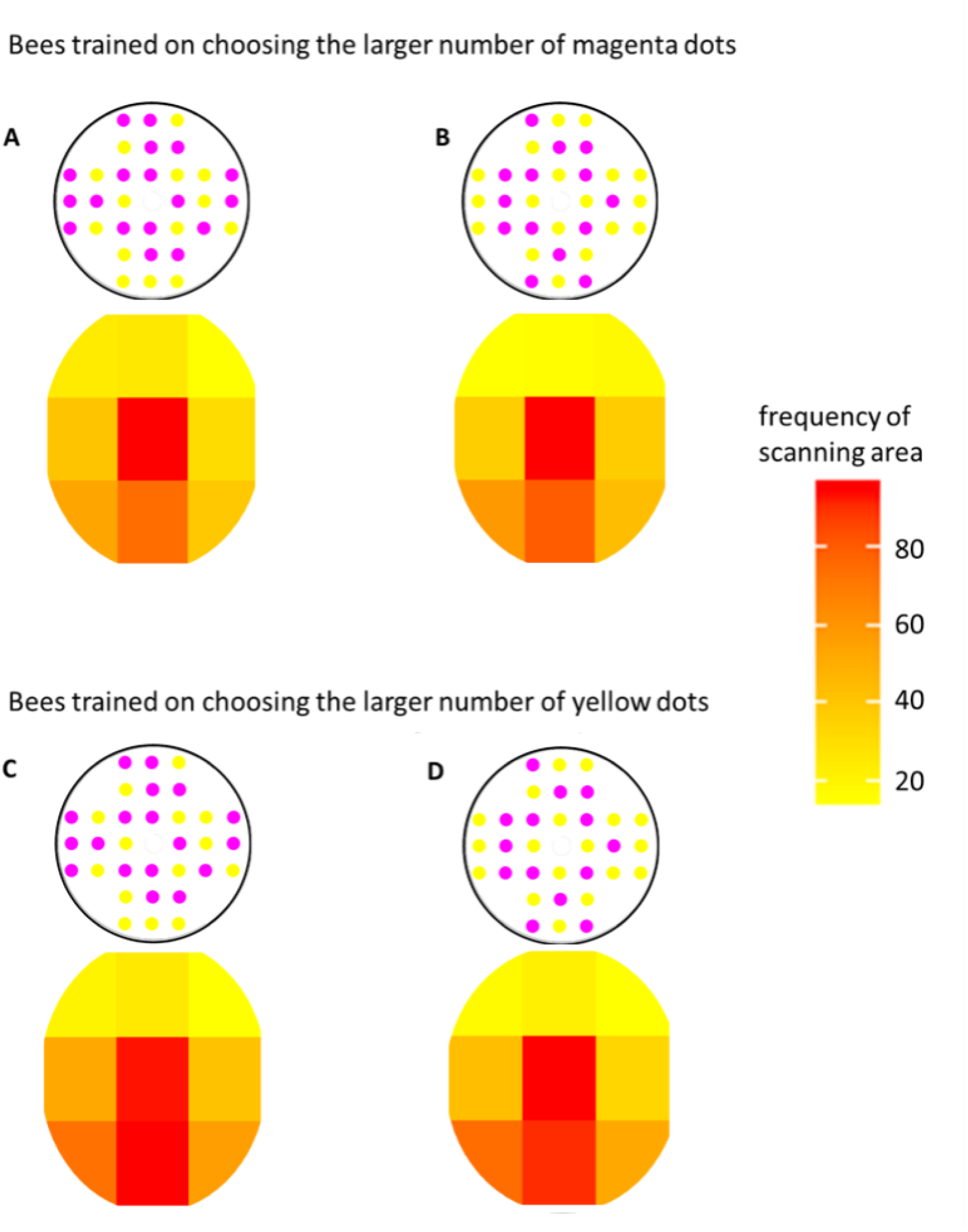
Scanning behaviour of bees in the presence of the generalization test. Frequency maps of bee location in front of a pattern during solitaire illusion tests. Bees spent significantly more time on scanning the lower and central areas of each pattern (bottom|middle: 95% CI [−0.51, −0.02]; middle|top: 95% CI [1.53, 2.06]).

In the “illusion test”, bumblebees showed a preference for the stimulus with clustered positive reinforced coloured as expected if they perceive the Solitaire Illusion. However, before claim that bumblebees make numerical judgment at glance, as humans do, or whether they might have showed similar preference but adopting different strategies, we focused on the scanning behaviour of bees. Bumblebees scanned the central area more frequently compared to the remaining areas of the pattern (see Figure 5). No significant difference was found between the types of configurations (cluster vs. spaced configuration: ß = 0.09, C.I. [−0.18, 0.34], χ^2^ = 0.00, *P* = 0.98). Neither the difference between the rewarded colour dots (ß = 0.02, C.I. [−0.27, 0.33], χ^2^_1_ = 0.31, *P* = 0.58), nor the interaction between these two factors was significant (ß = −0.17, C.I. [−0.59, 0.15], χ^2^_1_ = 0.98, *P* = 0.32; *McFadden’s* - *R*^2^ < 0.001). Analysis of hovering time revealed a significant effect of area (*F*_8,1621.03_ = 19.97, *P* < 0.01, ηp^2^ = 0.90), a significant effect of configuration (*F*_1,1631.98_ = 7.11, *P* < 0.01, ηp^2^ < 0.01), and, more importantly, a significant triple interaction area × configuration × rewarded coloured dots (*F*_8,1619.71_ = 3.77, *P* < 0.01, ηp^2^ = 0.02; Figure 6), suggesting that bumblebees scanned the central area of stimulus with clustered reinforced coloured dots for longer. No other factors were significant (*Ps* > 0.13). Considering the sequential scanned area during the flight path, the bumblebees’ choice has clearly been affected by the clustered configuration respect to the reinforced coloured dots (C.I. [0.75, 1.91], MCMC*P* < 0.01); bumblebees followed the line direction form by the clustered reinforced colour dots, especially the lower areas, and avoided the spared area of stimulus, suggesting that, even thought the behavioural outcome is the same between bees and humans, bees might not make numerical judgement at glance as human does, but they might use an alternative strategies such as following the spatial arrangement of reinforced colour dots.

**Figure 5.**
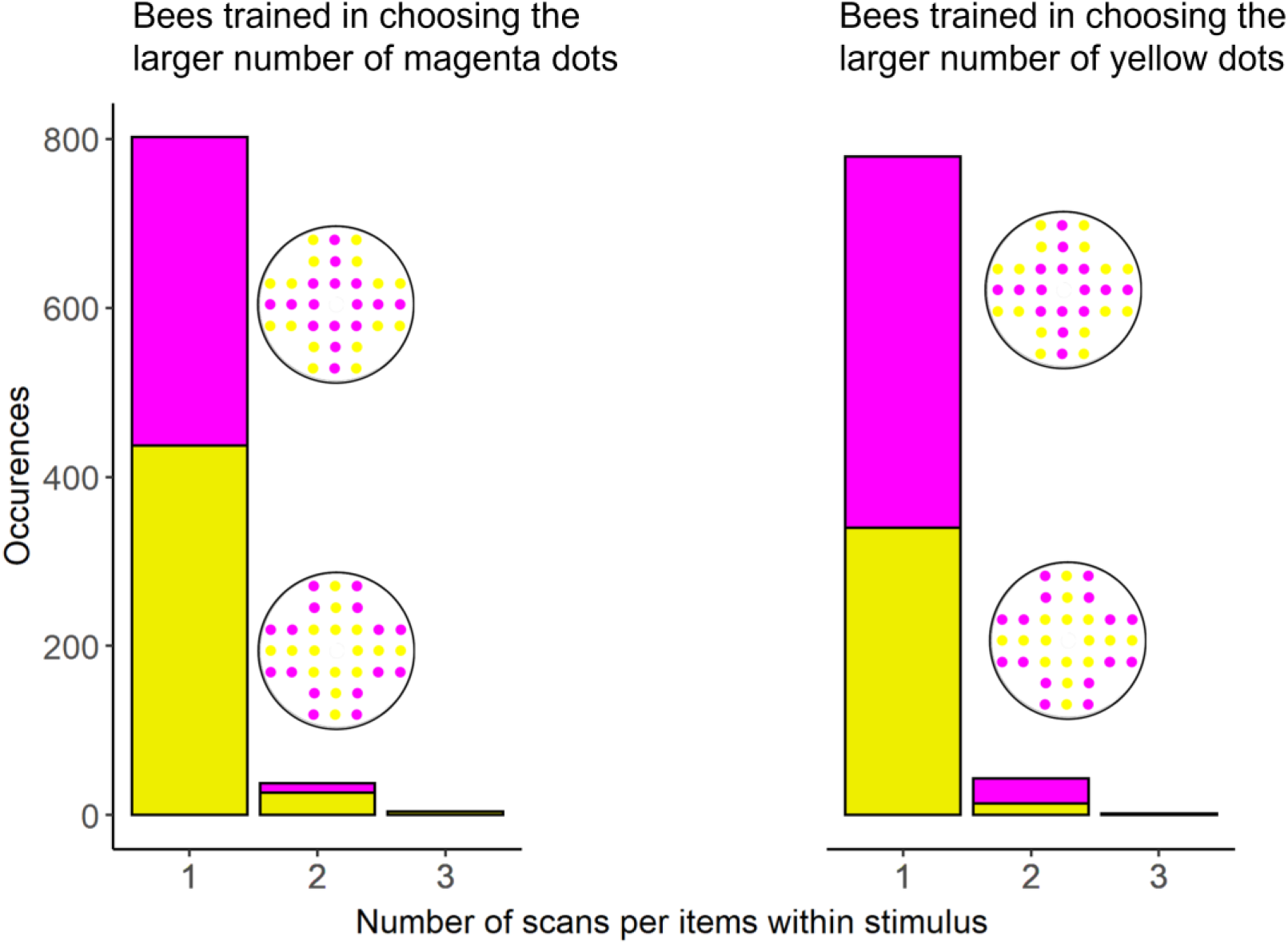
Sequential scanning of stimulus area by bumblebees. Cumulative number of occurrences of scanning behaviour for each area within stimulus. The left graph represented the scanning events of bees trained in choosing the larger number of magenta dots divided by the type of ‘numerosity illusion’ stimuli. The right graph represented the visit by bees trained in choosing the larger number of yellow dots.

**Figure 6.**
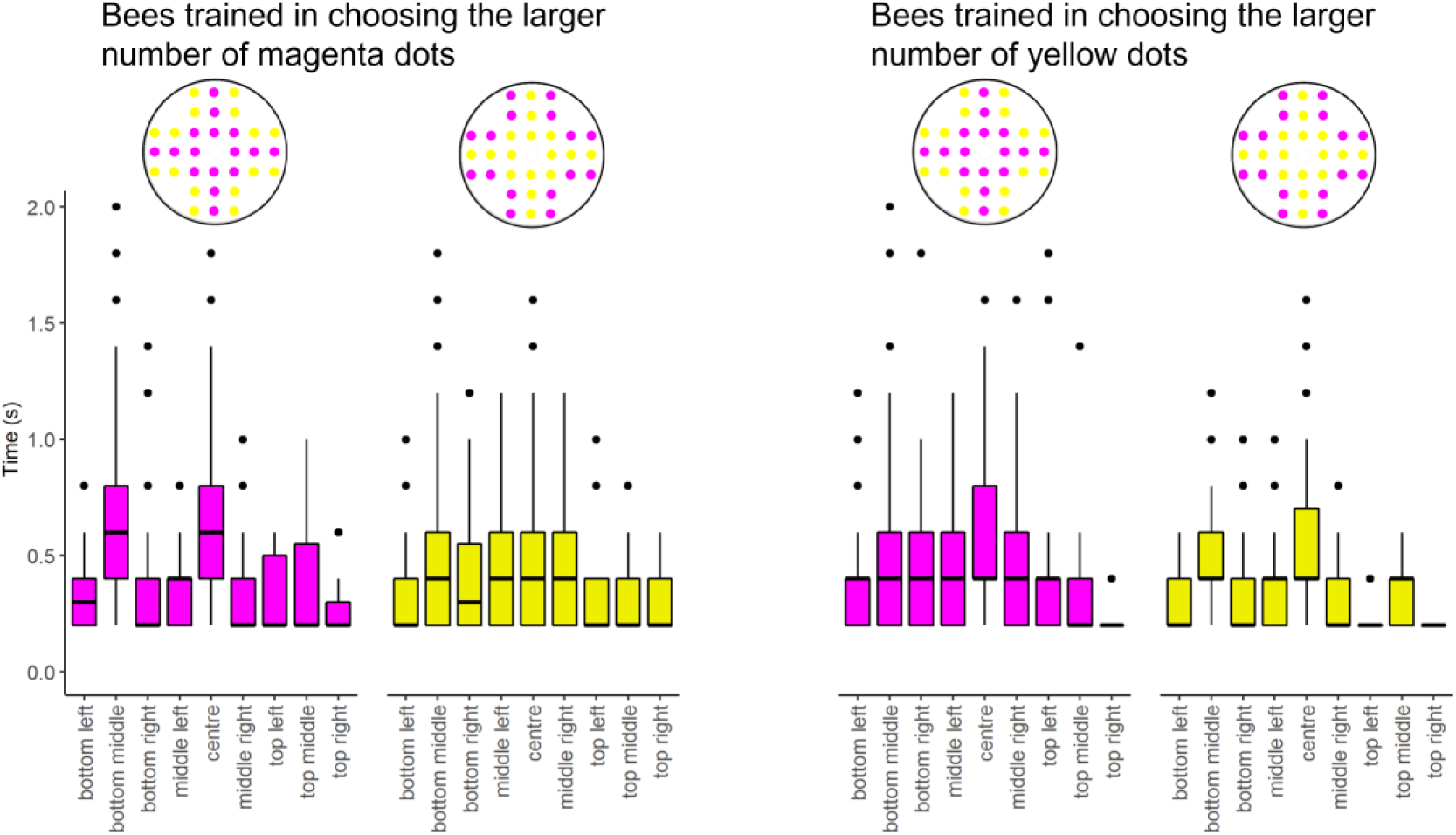
Time inspection of bees within each area of the Solitaire Illusion stimuli. The median (±SD) of time inspection plotted as a function of configuration pattern and the reinforced patterns. Overall, there is a significant triple interaction area × configuration × rewarded coloured dots (*P* < 0.01), suggesting that bees scanned longer the aggregation of reinforced colour dots compare to the other part of stimulus.

## Discussion

Despite a wide range of similarities in quantitative abilities of vertebrates [11–13, 30], the misperception of Solitaire illusions has been observed only in humans [9]. Here, we provided evidence that a species of bumblebee behaves as if it perceives the Solitaire Illusion (9) when only their final choices are observed. However, flying path analysis showed that bumbles might achieve the same behavioural results but not doing numerical judgement at glance as human does, but by scanning specifically the clustered coloured dots of the spatial configuration of Solitaire Illusion (9). These results raise intriguing questions about the mechanisms underlying visual processing.

A critical issue in visual perception studies is how different species integrate sensory information captured by the retina into the brain. A plethora of species seem to prioritize global structures of the elements rather than individual analysis of local properties, typically referred to as seeing ‘the forest before the trees’ [31-34 but see 35]. For example, a set of trees, which is closely spaced and similar to each other, lead them to be visually grouped into a forest. A forest has certain properties, i.e., density of elements, defined as Emergent Features, which emerge only when single parts are combined into a group [14]. The global component of an object concerns the spatial relationship between elements, while the local component regards the specific properties of single elements. According to the global precedence hypothesis [31], a visual object is represented from the global to local properties. Thus, the misperception of numerosity in the Solitaire Illusion, as well as other type of illusion [14], might be elicited by the spatial arrangement of elements (global properties) that influence the assessment of numerosity (local information) in each pattern. In our study, bumblebees showed a preference for clustered pattern as if they perceive the illusion, suggesting that bumblebees prioritize a global analysis of the stimuli rather than the local information of single elements. This hypothesis is consistent with previous proposals that global perception makes honeybee foraging more robust to disturbance of movement or viewing angle [36, 37].

However, before claiming that bumblebees perceive the illusion, we should consider alternative non-numerical explanation to describe our results. Indeed, video analyses showed that bees significantly scanned local elements, especially the area occupied by the centrally clustered reinforced colour dots, where the majority of same-coloured dots are allocated, and their spatial arrangement might guide bee to the center of stimulus. A recent study investigated the scanning strategies of bumblebees in discriminating between quantities [22]. Bumblebees were trained to distinguish patterns with two items from four and subsequently transfer numerical information to a novel numerical task. The video analyses of scanning behaviours revealed that bumblebees did not assess the number of elements at glance as the parallel process of visual search in mammals and birds; instead, bees sequentially scanned the countable items, similar to the motor tagging behaviours in primates, which consist of a prolonged visual search and store the temporal information in the working memory before making a decision. Although it was possible that bumblebees counted the amounts of coloured dots before making a numerical discrimination, our data might be interpreted in a different way. In the “illusion test”, the clustered dots formed a single continuous figure that led to the centre of stimulus from any position. Our video analysis supported this; bumblebees tend to sequentially follow the forming paths of the same reinforced colour dots when displayed in a clustered configuration; while no flight path emerged when the reinforced colour dots were peripherally located and separated by the dots of the opposite colour. Even thought previous studies showed that bumblebees might use numerical information [21, 22, 38], subjects might learn an alternative strategy (i.e., follow the line direction of same-colour dots) to solve the task.

In this study, we also found that an invertebrate species can discriminate between large quantities. A previous study showed that honeybees (*Apis mellifera*) can discriminate between 3 and 4 dots (ratio 0.75) in a delayed matching-to-sample task [38], cuttlefish (*Sepia officinalis*) between 4 and 5 preys (ratio 0.80) in a spontaneous food-choice task [39], and house crickets (*Acheta domesticus*) discriminate between 2 and 3 holes (ratio 0.67) when searching for potential refuges [40]. Not only did bees in our experiments learn to discriminate 11 vs. 21 dots (ratio 0.52) and one 13 vs. 19 dots (ratio 0.68), but also, they spontaneously transferred the learned rule to a novel discrimination, namely 14 vs. 18 (ratio 0.78). Our study does not contain direct evidence, however, that these large number discriminations are guided by counting the items in the various displays. Indeed, bees’ choices might be driven by non-numerical cues that correlate with numerosity. A larger group of objects usually had a larger cumulative surface area compared to a smaller group [41–43]. A recent study showed that some of the seemingly more advanced performances in counting tasks by bees, and possibly in other species, could be equally explained by the fact that individuals learned alternative strategies [44]. In a series of behavioural experiments, the authors reproduced several visual discrimination tasks commonly used to assess numerical cognition in bees. In this study, bees made magnitude-related decisions based on non-numerical information. Regarding our study, bumblebees could learn to search for clustered items with a larger cumulative area, or applying a rule based on the spatial frequency of elements (i.e. the repeating alternation of coloured dots in a pattern per unit distance), or following the line direction formed by the arrangement of elements, as likely shown in the ‘illusion test’. We cannot exclude this possibility when magenta and yellow dots differing in number were presented.

An interesting result emerges when we consider the sampling time taken by subjects before making a numerical discrimination. Bumblebees scanned the stimulus with smaller number of reinforced colour dots longer compared to the stimulus with larger number of reinforced colour dots. This behaviour can be interpreted as evidence that gathering accurate information in large-number displays required longer sampling time when performing challenging discrimination. Several studies reported that human and non-human animals showed a speed-accuracy trade-off in various decision-making contexts [45–47]. Indeed, we expected that bumblebees take longer sampling time to gather information when presenting with larger number discrimination before choosing/avoiding the stimulus [48–50]. Curiously in our study, bumblebees increased their sampling time when faced stimulus with smaller number of reinforced colour dots. This is counter to the results in small-number discriminations, where bees take longer to “tag” larger quantities of items [22], making it likely that bees used a non-numerical property to evaluate the displays in the present study.

## Materials and Methods

Experiments were performed with three colonies of *Bombus terrestris audax* (Biobest Belgium N.V., Westerlo, Belgium). Bees were maintained at Bee Sensory and Behavioural Ecology Lab, School of Biological & Chemical Sciences, Queen Mary University of London (UK) between December 2018 and February 2019. Each colony was connected to a wooden flight arena (100×70×70 cm) via a plastic tunnel. The arena was covered with UV-transparent Plexiglass ceiling.

Before starting the experiment, forager bees were attracted to the arena with white circular stimuli (Ø 12 cm, surrounded by 2 mm wide black margins) positioned on the vertical wall of the arena. Each white stimulus presented a feeder containing 30% sucrose solution. Individual forager bees were marked on their thorax with paint for identification during the experiment.

Stimuli (Figure 1B) consisted of arrays of magenta (RGB: 255, 0, 255) and yellow dots (RBG: 255, 255, 0; Ø 1 cm each dot) arranged in a cross-shape pattern (inter-object distance: 1 cm) on a white circular background (Ø 12 cm, surrounded by 2 mm wide black margins) made with Microsoft PowerPoint. We created four different configurations for each numerical comparison. Stimuli were laminated to allow cleaning with ethanol after each trial. All stimuli were printed with a high-resolution laser printer. Stimuli, feeding tubes, quinine and sugar solutions were replaced between each trial.

*Protocol*. In each experimental phase, four pairs of stimuli consisting of 32 dots were randomly placed on the vertical wall of the arena. The number of magenta and yellow dots within each pair was the same but with an inverse relation of one another with regard to specific quantities of each dot type. However, the number of magenta and yellow dots within each array varied according to the experimental phase. During the training phase bees were presented with 11 vs. 21 dots (ratio 0.52) and one 13 vs. 19 dots (ratio 0.68). During the learning test we presented again 13 vs. 19 dots (ratio 0.68), whereas in the generalization test bees were presented with 14 versus 18 dots (0.78 ratio). The correct and incorrect patterns were randomly assigned among subjects. Choosing the correct pattern was positively reinforced by 10 μL droplet of 50 weight % sucrose solution placed at the centre of the target stimulus inside a plastic feeder, while the incorrect pattern was negatively reinforced by 10 μL saturated quinine HCl solution.

During the training phase (Figure 1A), we administered nine trials where a single bee could freely forage in the arena. A trial was defined as a subject leaving the hive, choosing different stimuli, and freely returning to the nest once satiated. In our experimental protocol, a trial lasted on average 2 minutes (unpublished data). Since bees had not limitation of foraging time, the number of choices (e.g., landing and feeding in a stimulus) varied across subjects (total number of choices per subjects: 138.8 ± 40.39; average number of choices per training trial: 15.50 ± 3.52). During each training trial, empty positive and negative feeders were refilled after the bee had left the stimulus. We manually recorded all choices made by subjects in each trial. To assess learning performance, we considered the last 50 choices grouped as 5 group of 10 choices each. All selected foragers have successfully completed the training phase and no subjects were discarded.

After the training, we administered three tests divided by two refreshing training trials to maintain subjects highly motivated on foraging. In all the test phases, the stimuli provided 10 μL of sterilized water. We recorded the behaviour for 2 min with a camera positioned at the entrance of the arena. During the test trial, the experimenter manually collected all the correct and incorrect choices made by subjects to assess their performance. The recorded videos were subsequently analysed using the free software Solomon coder beta (Andras Peter, BUDAPEST). In the learning and the generalization tests, we firstly measured the hovering time (i.e., the time spent hovering in front of a pattern) for each visit. Then, we virtually divided each stimulus in three equal virtual area, defined as bottom, middle, and top according to the vertical axis of the stimulus, to evaluate which area were more informative for the bees before making a decision. For the “illusorion test”, we further analysed the scanning behaviour by dividing in nine equal area according to the spatial arrangement of the two-coloured arrays. For each virtual area, we measured the number of visit and the hovering time.

An additional group of bees (n = 10) were tested in a control experiment to evaluate spontaneous preference for a spatial configuration. Bees were presented with two different arrays: one array presented centrally located dots, the other presented dots located in the perimeter to form four separate clusters. Pairs of stimuli were made with the same number of magenta or yellow dots. We tested 5 bees for each pair of stimuli as described above.

One may argue that the significant choice observed in the illusory trials may not be the result of a genuine misperception of numerosity, but instead reflect a spontaneous preference for the visual patterns in which the target dots were centrally located (a pattern that for instance might resemble a flower). To test this hypothesis, we set up a control experiment (n = 10) in which bees were observed in their natural tendency to explore two different arrays: one array presented centrally located dots, the other presented dots located in the perimeter to form four separate clusters. Bees did not explore one array more than chance (performance 51.63% ± 18.33; *Estimate* = 0.137, *SD* = 0.214, *z* = 0.639, *P* = 0.523; effects of colour dots: χ^2^_1_ = 0.076, *P* = 0.783, *delta - R*^2^ < 0.001), showing that the choice for the arrays with centrally located target dots in the training study is not due to any form of a-priori preference for a given array.

## Statistical analysis

Statistical analysis was performed in R version 3.2.0 (The R Foundation for Statistical Computing, http://www.r-project.org). For the training phase and test phase, we analysed the choices (correct or incorrect) with generalized linear mixed-effects models for binomial response distributions (GLMMs, ‘glmer’ function of the ‘lme4’ R package). To evaluate the performance during the training phase, the GLMM model was fitted with block number (10 consecutive choices) as covariate (to establish whether performance increased over trials), the positive reinforced colour as factor (to assess whether performance differed for the reinforced pattern), individual ID as a random effect, and log as link function. To evaluate the performance in the test phases, the GLMM fitted with reinforced pattern as factor, individual ID as a random effect, and logit as link function. To assess the significance of the models’ parameters, we used the ‘Anova’ function of the ‘CAR’ package.

In the learning and transfer tests, we analysed the scanning behaviour from video-recordings in two steps. The first step was focused on evaluating difference in time inspection when bumblebees scanned both patterns. We analysed the hovering time by using a GLMM model with Gaussian distribution family (i.e., Linear Mixed-Model, LMM) fitted with the positively rewarded pattern and the configuration of array as fixed factors, and individual ID as random effect. The second step was focused evaluating scanned area of stimuli. We analysed the scanned area of each pattern (i.e., bottom, middle, and top for the learning and generalization test; the nine equal sectors for the illusory test) with a cumulative link mixed-effects models (CLMM, ‘clmm’ function of the ‘ordinal’ R package) fitted with the positively rewarded pattern and the configuration of array as fixed factors, and individual ID as random effect. To assess the significance of the models’ parameters, we used the ‘Anova’ function of the ‘RVAideMemoire’ package. We reported the pseudo-R2 (i.e., McFadden’s - R2) by comparing the null model without fixed effects except for an intercept and the full model with the ‘nagelkerke’ function of the ‘rcompanion’ package.

A similar approach was applied for analysing the scanning behaviour during the “illusion test”. Firstly, we analysed the scanned area of each pattern (i.e., nine different virtual area) by using a CLMM models fitted with the positively rewarded pattern and the configuration of array as fixed factors, and individual ID as random effect. Secondly, we analysed the hovering time with a LMM fitted with the virtual area, the positively rewarded pattern, and the configuration of array as fixed factors, and individual ID as random effect. We used a Bayesian generalized linear mixed models to estimate the probability of bees to sequentially searching for similar pattern. We analysed bees’ choice by using a MCMCglmm model (‘MCMCglmm’ function in ‘MCMCglmm’ R package, [51]) fitted with type of configuration and reinforced colour as fixed factors, and individual ID as random effect. We considered a non-informative prior (V = 1, nu = 0.002, α.mu = 0, α.V = 1000, [52]) and ran the model for 14000 with a burn-in of 4000 that produced 1000 estimates of the posterior distribution.

## Ethics

Italian and United Kingdom law do not require ethical approval for studies involving insects. However, all animals in our study were maintained and tested following the ASAB/ABS Guidelines for the Use of Animals in Research [51].

## Acknowledgments

We would like to thank Luigi Baciadonna for insightful discussions. This research was financed by ERC grant “SpaceRadarPollinator” (grant code: 339347) by L.C., Horizon Europe Framework Programme grant NimbleAI by L.C, PRIN 2015 Grant (Prot. 2015FFATB7) from University of Padova by A.C., and FAR2022 (2022-FAR.L-GE_001) from University of Ferrara by E.G.

## Author contributions

All the authors equally conceived the study and contributed to the final version of the manuscript; EG and CG collected the data; EG analysed the data; EG, AC, LC, and MEMP drafted the manuscript.

## Data availability statement

The data that support the findings of this study are available from the corresponding author, E.G., upon reasonable request.

---

**Video S1 Example video of the flight path during the solitaire illusion test of a subjects previously trained to select patterns with larger number of magenta dots.** The bee sequentially examined items of the patterns containing yellow dots in a clustered configuration but rejected it after scanning. Later, bees scanned and landed on a pattern with magenta dots in a clustered configuration. At the end, bees scanned and rejected another pattern with yellows dots in a clustered configuration.

